# Identification and characterization of yeast and human glycosphingolipid flippases

**DOI:** 10.1101/373712

**Authors:** Bartholomew P. Roland, Tomoki Naito, Jordan T. Best, Cayetana Arnaiz-Yépez, Hiroyuki Takatsu, Roger J. Yu, Hye-Won Shin, Todd R. Graham

**Author notes:** These authors contributed equally to this work (Bartholomew P. Roland, Tomoki Naito). Correspondence: Todd R. Graham.

## Abstract

Lipid transport is an essential process with manifest importance to human health and disease. Phospholipid flippases (P4-ATPases) transport lipids across the membrane bilayer, and are involved in signal transduction, cell division, and vesicular transport. Mutations in flippase genes cause or contribute to a host of diseases such as cholestasis, neurological deficits, immunological dysfunction, and metabolic disease. Genome-wide association studies have shown that *ATP10A* and *ATP10D* variants are associated with an increased risk of diabetes, obesity, myocardial infarction, and atherosclerosis; and *ATP10D* SNPs are associated with elevated levels of glucosylceramide (GlcCer) in plasma from diverse European populations. Although sphingolipids are strong contributors to metabolic disease, little is known about how GlcCer is transported across cell membranes. We have identified an evolutionarily conserved clade of P4-ATPases from *Saccharomyces cerevisiae* (Dnf1, Dnf2), *Schizosaccharomyces pombe* (Dnf2), and *Homo sapiens* (ATP10A, ATP10D) that transport GlcCer. Further, we establish the structural determinants necessary for the specific recognition of this sphingolipid substrate. Our molecular observations clarify the relationship between these flippases and human disease, and have fundamental implications for membrane organization and sphingolipid homeostasis.

## Introduction

Phospholipid flippases in the P4-ATPase family establish and repair the phospholipid asymmetry of cellular membranes, and facilitate import of phospholipid from extracellular sources (Roland & Graham, 2016a). These enzymes are integral membrane proteins that harness energy from ATP catalysis to translocate lipids from the exofacial leaflet to the cytofacial leaflet of cellular membranes. There are 14 distinct P4-ATPases in humans with important differences in tissue-specific expression, subcellular localization, and substrate specificity (Andersen, Vestergaard et al., 2016). These differences in P4-ATPase localization and enzymology are important for coordinating physiological processes, and their impairment results in diverse pathologies (Folmer, Elferink et al., 2009). For example, the P4-ATPase ATP8B1 transports phosphatidylcholine at the apical membrane of canalicular hepatocytes (Takatsu, Tanaka et al., 2014), and mutations in the *ATP8B1* gene lead to membrane damage and hepatic cholestasis (Bull, van Eijk et al., 1998, Folmer et al., 2009). Conversely, ATP8A2 translocates phosphatidylserine within the central nervous system, and mutations in *ATP8A2* cause cerebellar ataxia mental retardation and disequilibrium syndrome (CAMRQ) in humans, and motor neuron degeneration in mice (Coleman, Zhu et al., 2014, Onat, Gulsuner et al., 2013, Zhu, Libby et al., 2012). Thus, defining the substrate specificity of the P4-ATPases is crucial to understanding their role in disease.

Genome-wide association (GWA) studies have found that single nucleotide polymorphisms (SNPs) in the human P4-ATPase genes *ATP10A* and *ATP10D* are strongly linked with metabolic disease. One GWA study of non-diabetic African American patients identified two SNPs in *ATP10A* that associated with insulin resistance (Irvin, Wineinger et al., 2011), while two GWA studies of European and Japanese populations have revealed strong associations between commonly found *ATP10D* SNPs and both myocardial infarction and atherosclerosis (Hicks, Pramstaller et al., 2009, Kengia, Ko et al., 2013). Importantly, analyses of plasma sphingolipids from over 4000 individuals revealed a specific elevation in glucosylceramide (GlcCer) levels associated with *ATP10D* SNPs (Hicks et al., 2009).

Mouse models with *Atp10a* and *Atp10d* mutations also demonstrate a link to metabolic disease. Two mouse strains with *Atp10a* mutations (formerly known as *Atp10c*) are susceptible to diet-induced insulin resistance and obesity (Dhar, Hauser et al., 2002, Dhar, Sommardahl et al., 2004). Additionally, the C57BL/6 mouse strain, which carries a premature stop codon in an *Atp10d* exon (Flamant, Pescher et al., 2003), is specifically prone to develop obesity, hyperglycemia, and hyperinsulinemia under a high-fat diet (Surwit, Feinglos et al., 1995); and these metabolic phenotypes can be attenuated by the transgenic complementation of *Atp10d* (Sigruener, Wolfrum et al., 2017). Lipidomic examinations of *Atp10d*-deficient mice demonstrated that GlcCer was elevated in mouse plasma, similar to the human patients, and that complementation with the *Atp10d* transgene also mitigated this increase (Sigruener et al., 2017).

No substrates have been reported for the enzyme ATP10D. P4-ATPases were presumed to selectively translocate glycerophospholipids and not sphingolipids; therefore, the molecular basis for the links between *ATP10D* and GlcCer was mysterious. In addition, the ATP10D-related P4-ATPases in *Saccharomyces cerevisiae* (Dnf1 and Dnf2) are part of a regulatory circuit that controls sphingolipid synthesis through an as-yet-unclear mechanism (Roelants, Baltz et al., 2010). Sphingolipids are divided into four classes: i) sphingoid bases, ii) ceramides, iii), phospho-sphingolipids, and iv) glycosphingolipids (Vance & Vance, 2008). GlcCer is the central metabolite between two of these major classes, the ceramides and glycosphingolipids. GlcCer is one commonly found sphingolipid in the circulation, though its transport to and from target tissues is poorly understood. Like ceramide, GlcCer is carried through the body by lipoproteins (Dawson, Kruski et al., 1976), though its amphipathic chemistry suggests that it requires active transport for absorption and distribution across the membranes of target tissues. Identifying and characterizing GlcCer transporters will be essential for testing the systemic and tissue-specific roles of this lipid in metabolic disease.

In previous studies, we and others have examined the molecular determinants of P4-ATPase substrate specificity (Baldridge & Graham, 2012, Stone, Chau et al., 2012, Vestergaard, Coleman et al., 2014). The yeast P4-ATPases in particular have been indispensable for understanding how phospholipid headgroups and backbones are coordinated/selected during transport. In this study, we have identified a conserved clade of yeast and human P4-ATPases that transport GlcCer. Further, a central glutamine in TM4 is conserved in all identified GlcCer-flippases, and is necessary for GlcCer but not glycerophospholipid transport. Identifying and characterizing these transporters significantly advances our understanding of systemic lipid delivery in mammals, and provides the enzymological basis for targeted pharmaceutical development. We propose that these enzymes function as critical sphingolipid importers within fungi and humans, and that GlcCer transport likely evolved to support cellular lipid homeostasis.

## Results

### Fungal P4-ATPases transport glycosphingolipids

Dnf1 and Dnf2 are two related P4-ATPases that localize to the plasma membrane of *S. cerevisiae*, and transport phosphatidylcholine (PC) and phosphatidylethanolamine (PE). To probe how P4-ATPases distinguish glycerophospholipids from sphingolipids, we previously mutagenized the budding yeast P4-ATPase Dnf1 and selected variants capable of transporting sphingomyelin (SM). Gain-of-function mutations were identified (Dnf1^N200S,L1202P^) that changed the substrate preference of Dnf1 from PC/PE to SM (Roland & Graham, 2016b). While testing the specificity of these SM-permissive Dnf1 mutants, we surprisingly discovered an existing capacity for wild type (WT) Dnf1 to transport GlcCer and galactosylceramide (GalCer) (Fig. S1). Therefore, we broadly examined the requirement for Dnf1 and Dnf2 in sphingolipid transport relative to known substrates using WT; *dnf1∆*; *dnf2∆*; and *dnf1,2∆* knockout (KO) strains. As previously reported, (7-nitro-2-1,3-benzoxadiazol-4-yl)-PC (NBD-PC) and NBD-PE transport progressively decreased in each KO strain (Fig. 1a, Fig. S2c) (Pomorski, Lombardi et al., 2003). WT *S. cerevisiae* surprisingly transported substantially more NBD-GlcCer and GalCer than NBD-PC, which was thought to be the preferred substrate for Dnf1 and Dnf2. Dnf2 is the primary NBD-GlcCer transporter at the plasma membrane because *dnf2∆* caused the greatest reduction in transport relative to WT cells. After uptake, NBD-GlcCer accumulated in the mitochondria (Fig. 1b) and was stable for up to 2 hours after administration (Fig. S3). NBD-GlcCer was not found to accumulate in the ER or endosomes regardless of P4-ATPase expression (Fig. S4). None of the P4-ATPase KO strains displayed a decrease in NBD-SM, NBD-ceramide (Cer), or NBD-lactosylceramide (LacCer) transport (Fig. 1a), suggesting that these lipids are not Dnf1 or Dnf2 substrates. These experiments demonstrate that the Dnf1 and Dnf2 flippases are capable of transporting monosaccharide-but not disaccharide glycosphingolipids.

**Fig 1.**
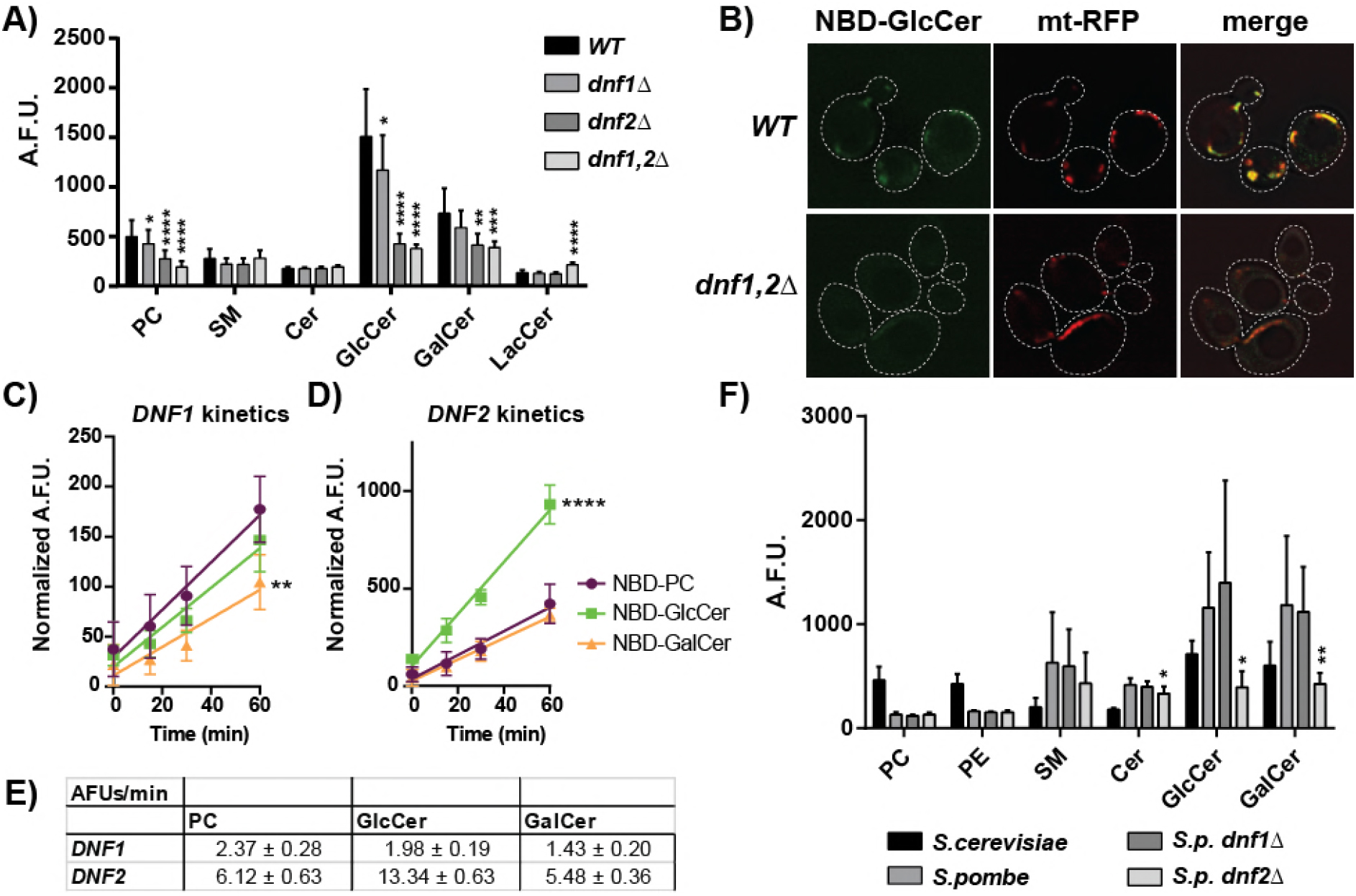
*S. cerevisiae* NBD-monosaccharide glycosphingolipid uptake requires plasma membrane P4-ATPases, Dnf1 and Dnf2. **(A)** NBD-lipid uptake was measured in WT (BY4741) and P4-ATPase knockout strains and presented as raw, arbitrary fluorescent units (A.F.U.), n≥9 ± SD. **(B)** Upon Dnf1,2-dependent uptake, NBD-GlcCer localized to mitochondria (mt-RFP). Kinetics assessments of NBD-PC, NBD-GlcCer, and NBD-GalCer uptake in *dnf1,2Δ* cells expressing pRS313-DNF1 (**C**) or pRS313-DNF2 (**D**), normalized to empty vector controls, n=6 ± SD. **(E)** Velocities of substrate transport from linear regression fits of data in panels C and D, ± S.E. **(F)** NBD-lipid uptake was measured in *S.c.* WT, *S.p.* WT, and *S.p. dnf1Δ* and *dnf2Δ* strains. Asterisks in panel F indicate differences in *S.p.* KO strains from *S.p.* WT. A one-way ANOVA was performed to assess variance in panels A and F, and comparisons made with Tukey’s post hoc analysis. A Two-way repeated measures ANOVA was used to assess variance in the kinetic data, and comparisons to NBD-PC were made with Tukey’s post hoc analysis: * indicates p < 0.05, ** p < 0.01, *** p < 0.001, and **** p < 0.0001.

The magnitude of NBD-GlcCer uptake suggested that it may be the preferred substrate of Dnf1 and Dnf2. Therefore, we measured the kinetics of NBD-PC, NBD-GlcCer, and NBD-GalCer transport (Fig. 1c-e). Plasmid-borne *DNF1* and *DNF2* constructs complemented lipid transport in a *dnf1,2∆* strain (Fig S2a), and were used to normalize the genetic background to empty vector controls. Dnf1 transported NBD-PC and NBD-GlcCer at equivalent rates, and NBD-GalCer to a lesser extent (Fig. 1c, e). Comparatively, Dnf2 transported NBD-GlcCer at twice the rate of NBD-PC and NBD-GalCer (Fig. 1d, e). Competition assays demonstrated that GlcCer lacking the N-acyl chain and NBD was capable of acutely inhibiting NBD-GlcCer uptake by Dnf2 (Fig S2d), implying that the enzyme is capable of recognizing unmodified glycosphingolipid.

We then examined a distantly related fungal species, *Schizosaccharomyces pombe*, to test the conservation of P4-ATPase-mediated GlcCer transport. Measurements of NBD-lipid transport in *S. pombe* revealed robust uptake of GlcCer and GalCer, surpassing that of WT *S. cerevisiae* (Fig. 1f and Fig S2b, c). Deleting the *S. pombe DNF2* ortholog elicited a substantial reduction in glycosphingolipid transport (Fig. 1f and Fig S2b, c). Surprisingly, *S.p.* Dnf2 did not transport NBD-PC or NBD-PE, indicating GlcCer/GalCer transport is the conserved function of this fungal Dnf2.

### Human P4-ATPases specifically transport glucosylceramide

Eight of the fourteen human P4-ATPases localize to the plasma membrane (Bryde, Hennrich et al., 2010, Takatsu, Baba et al., 2011, van der Velden, Wichers et al., 2010) and were tested for the ability to transport sphingolipids across the plasma membrane of HeLa cells (Fig. 2a-c, and Fig S5). No substrate had previously been reported for ATP10D, yet this flippase was capable of transporting NBD-GlcCer (Fig. 2a, c, and d) with exquisite specificity; NBD-GalCer was not transported (Fig. 2a). The ATP10D^E215Q^ mutant lacks ATPase activity, and its inability to transport NBD-GlcCer demonstrates that translocation requires ATP catalysis (Fig. 2b). ATP10A preferred NBD-PC as previously reported (Fig. 2c) (Naito, Takatsu et al., 2015, Takada, Naito et al., 2018), but measuring the kinetics of GlcCer uptake relative to parental (-) and ATP10D^E215Q^ controls revealed that ATP10A was also capable of transporting NBD-GlcCer, though at half the rate of ATP10D (Fig. 2d, e).

**Fig 2.**
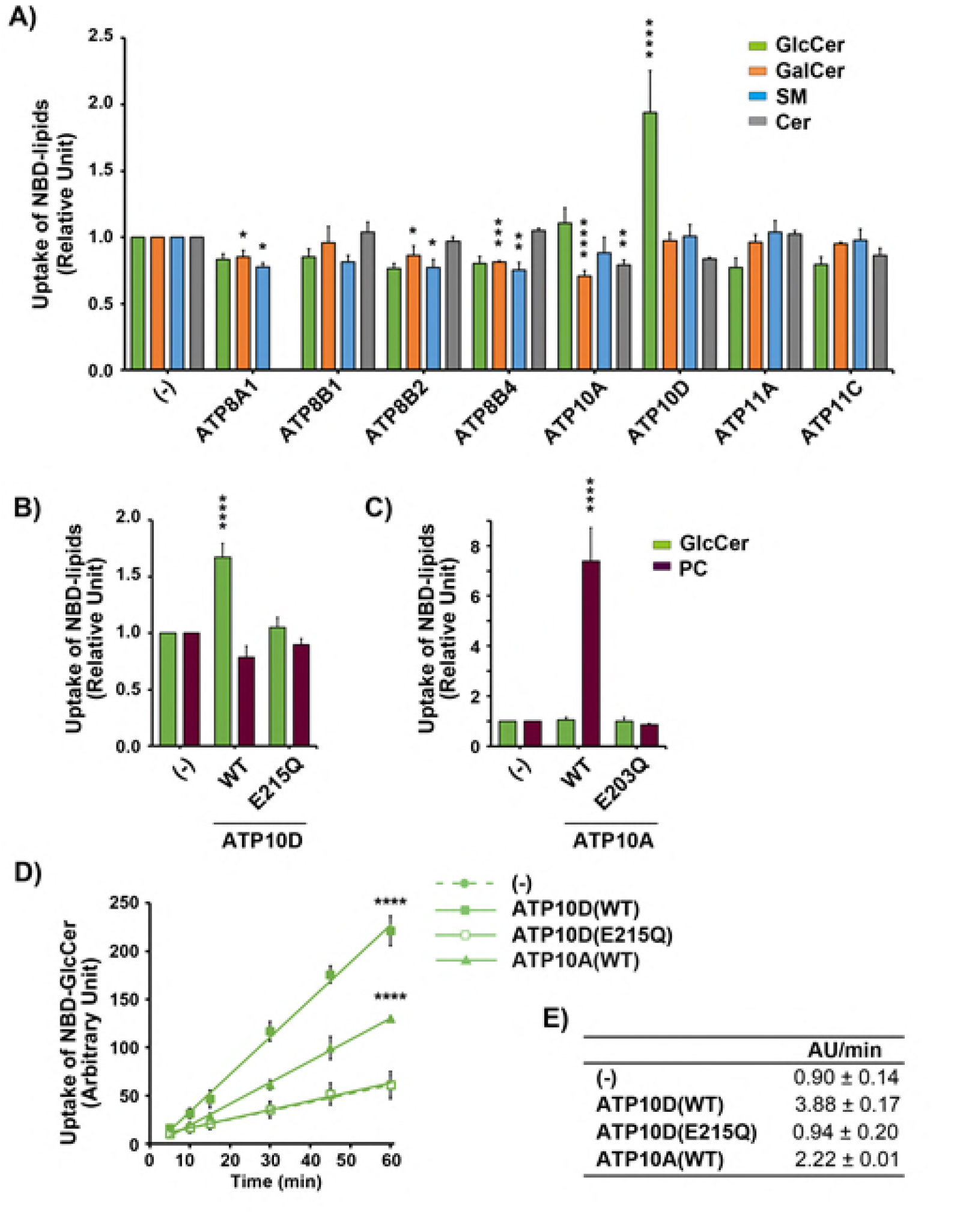
*H.s.* ATP10A and 10D translocate NBD-GlcCer at the plasma membrane. (**A**-**C**) Parental HeLa cells (-) and cells stably expressing human P4-ATPases and mutants were incubated with the indicated NBD-lipids at 15°C for 15 min. After extraction with fatty acid-free BSA, the residual fluorescence intensity associated with the cells was determined by flow cytometry. Graphs display averages from three to four independent experiments ± S.D. Fold increase of NBD-lipid uptake compared with parental cells (-) is shown. A one-way ANOVA was performed to assess variance in panels **A**-**C**, and comparisons to parental cells (-) were made with Tukey’s post hoc analysis. **(D)** HeLa cells were incubated with NBD-GlcCer at 15°C for indicated times (x axis). **(E)** Velocities of GlcCer transport from linear regression fits of data in panel D, ± S.D. Graphs display averages from three independent experiments ± S.D. A Two-way repeated measures ANOVA was used to assess variance in the kinetic data, and comparisons to (-) were made with Tukey’s post hoc analysis: * indicates p < 0.05, ** p < 0.01, *** p < 0.001, **** p < 0.0001.

### TM1 and TM4 participate in GlcCer transport

P4-ATPase substrate selection and transport are coordinated by the first six transmembrane (TM) domains, and chimeric mapping of these segments has been a productive strategy for dissecting P4-ATPase enzymology (Baldridge & Graham, 2012, Baldridge & Graham, 2013, Jensen, Costa et al., 2017). We used a series of chimeras between Dnf1 and Drs2 (a phosphatidylserine flippase (Natarajan, Wang et al., 2004)) to broadly map the transmembrane segments required for NBD-GlcCer transport (Baldridge & Graham, 2012). Dnf1 chimeras bearing Drs2 TM segments 1-4 displayed a significant reduction in NBD-GlcCer transport, and two-fold changes to the ratio of GlcCer to PC (a measure of substrate preference) (Fig S6). A focused examination of these regions revealed one motif in TM1 (GA) and two in TM4 (WVAV; YQS) that were particularly important for GlcCer preference (Fig S7). Two additional regions in the lumenal loop connecting TM1 and 2 (LL1-2) impaired both PC and GlcCer transport equivalently, and thus were considered general loss-of-function (Fig S7a).

Dnf2 is the primary PM-localized GlcCer transporter in *S. cerevisiae* (Fig. 1a, 1d) and its transport activity provides a greater dynamic range of responses. All three GlcCer motifs are conserved in the Dnf1 paralog, Dnf2 (Fig S7d); therefore, we performed a more thorough examination of these motifs in Dnf2. P4-ATPase homology models predict that the GA motif is positioned near the membrane/exoplasmic interface of TM1 (Fig. 4e). A sequence Logo and alignment illustrate the variability of this motif and the surrounding area (Fig. 3a, b). Substitutions at the glycine and alanine positions reduced the GlcCer/PC transport ratio but did not impact PC or PE transport (Fig. 3c, d). Double mutants in this region had the most profound impact on GlcCer preference and transport (Fig. 3e, f). We previously linked the QQ motif at this same position to phosphatidylserine selection in Drs2, a Golgi-resident P4-ATPase (Baldridge & Graham, 2013). Additionally, the exoplasmic region of TM1 was also involved in substrate coordination and transport in a human phosphatidylserine-translocating P4-ATPase, ATP8A2 (Gantzel, Mogensen et al., 2017), though the QQ motif was not directly examined. These studies collectively indicate that this GA/QQ motif in TM1 is an important component of GlcCer/PS recognition, but does not contribute to PE/PC selection.

**Fig 3.**
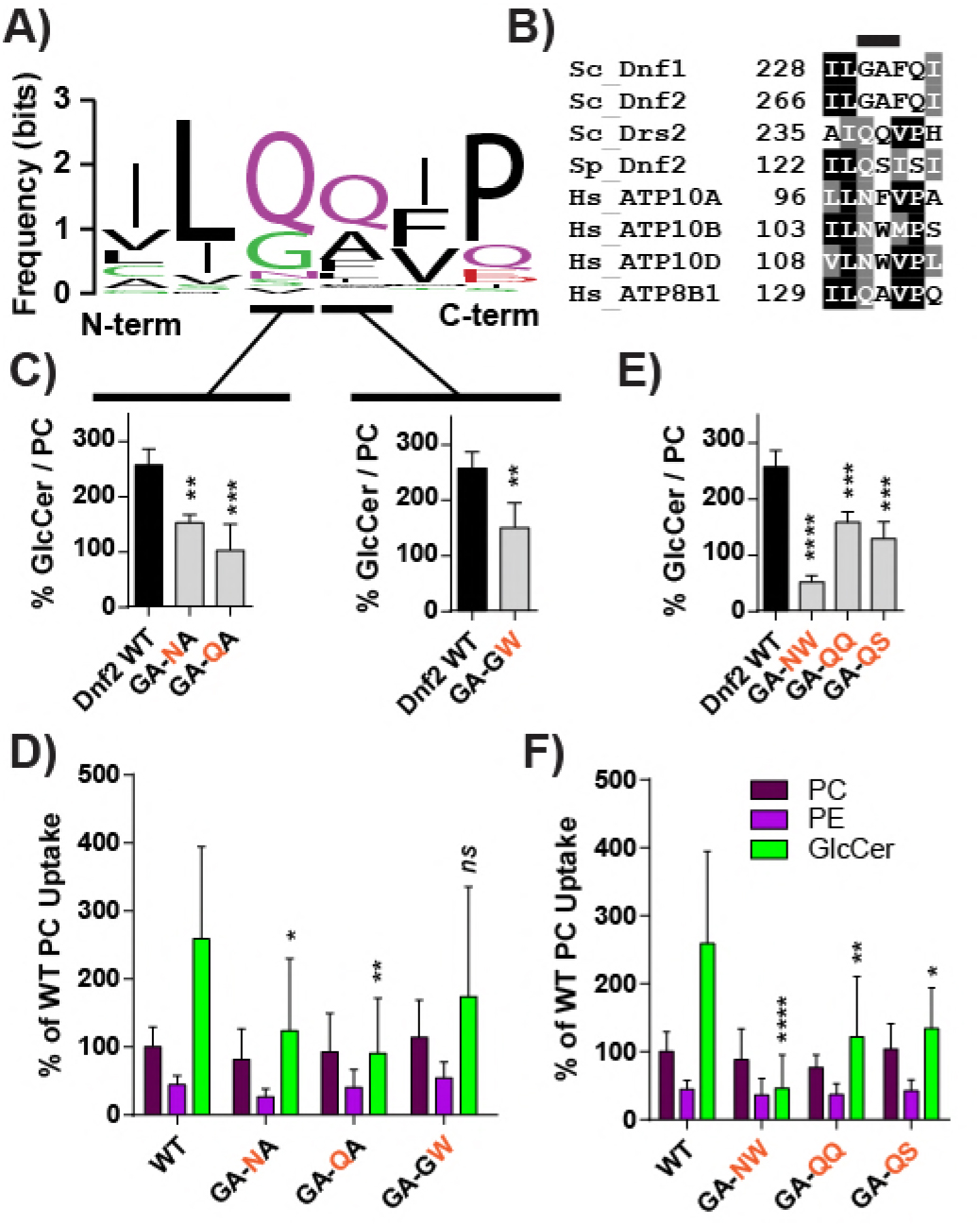
The exofacial TM1 “GA motif” facilitates Dnf2 selection and preference for GlcCer. **(A)** A sequence logo was created from an alignment of TM1 of P4-ATPases from different organisms, with letter size representing residue frequency and color denoting chemical differences. Hydrophilic residues are **green** and **purple**, acidic residues in **red**, and hydrophobic residues are indicated in **black**. **(B)** A focused alignment comparing a region of TM1 from *S.cerevisiae*, *H.sapiens*, and *S.pombe*, highlighting the **GA** motif. (**C-D**) Dissecting the first and second positions of the GA motif reveals that substitutions in both positions can reduce GlcCer preference (**C**), but do not alter PC or PE recognition (**D**). (**E-F**) Double-substitutions were created to examine *S. pombe* and *H. sapiens* sequences in the context of the *S. cerevisiae Dnf2*. These compound mutations reduced GlcCer preference (**E**) and selection (**F**) without altering the known glycerophospholipid substrates (**F**). Variance was assessed among data sets using One-way ANOVAs, and comparisons to WT made with Tukey’s post hoc analysis. Although first and second position GA analyses are presented in different panels (**C**), their statistical variance was tested together. * indicates p<0.05, **p<0.01, ***p<0.001, ****p<0.0001.

**Fig 4.**
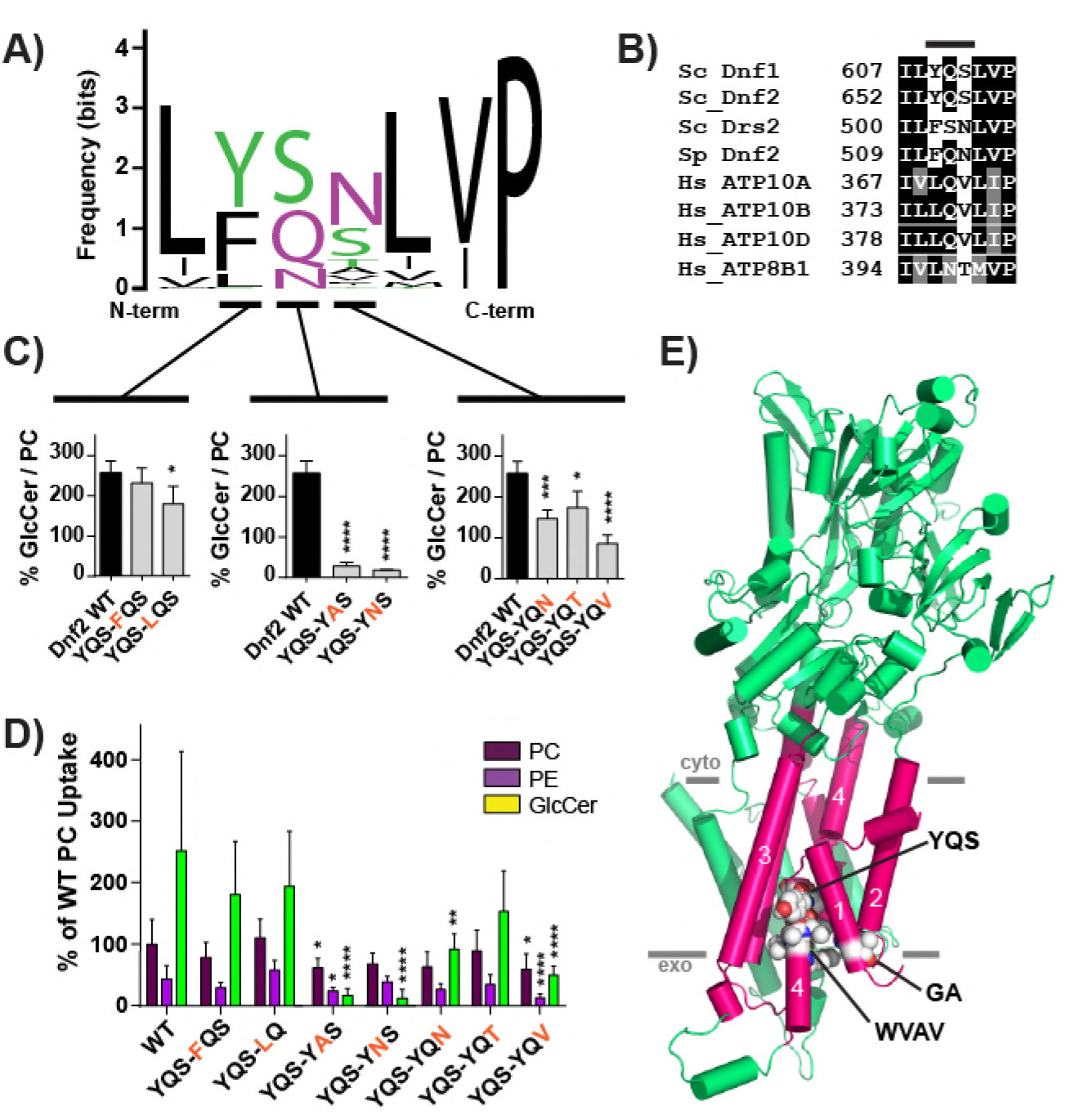
A Pro-4 glutamine in TM4 of Dnf2 is required for GlcCer transport. **(A)** A sequence logo was created from an alignment of TM4 of P4-ATPases from different organisms, with letter size representing residue frequency and color denoting chemical characteristics. Hydrophilic residues are **green** and **purple**, acidic residues in **red**, and hydrophobic residues are indicated in **black**. **(B)** A focused alignment comparing a region of TM4 from *S.cerevisiae*, *H.sapiens*, and *S.pombe*, highlighting the **YQS** motif that was previously altered in *S.c. Dnf1* (fig. S7). (**C-D**) The three **YQS** positions were tested for their influence on GlcCer preference (**C**) and selection (**D**), revealing that the central glutamine was the strongest determinant of GlcCer transport. (**E**) Homology model of Dnf1 with TM 1–6 shown as pink cylinders, the rest of the protein colored green. GA, YQS, and WVAV motifs are represented in spheres and colored by element. PM boundaries are indicated. Variance was assessed with one-way ANOVAs, and comparisons to WT made with Tukey’s post hoc analysis. Although the YQS positions in **C** are presented in separate panels, their statistical variance was tested together. n ≥ 9, ± SD, * indicates p<0.05, **p<0.01, ***p<0.001, ****p<0.0001.

TM4 is predicted to be a long alpha-helix, broken in the center by a conserved proline found in all P-type ATPases; the WVAV and YQS motifs are exoplasmic relative to this central proline (Fig. 4e). Replacing WVAV in Dnf1 with LTFW from Drs2 reduced PC transport and eliminated GlcCer selection (Fig S7b). However, single-position mutations in the WVAV motif of Dnf2 did not alter GlcCer preference or selection (Fig S8c, d), suggesting that the general integrity of this region may be more critical to lipid transport than individual amino acid side chains.

The YQS motif of TM4 is a functionally conserved island of polarity flanked by numerous hydrophobic residues (Fig. 4a, b). Y^654^ mutations modestly influenced GlcCer selection but changes to S^656^, including a Ser to Thr mutation, caused a more substantial and specific loss of GlcCer transport (Fig. 4c, d). However, altering the central glutamine (Dnf2^Q655A/N^) elicited the strongest perturbation of GlcCer preference and transport (Fig. 4c, d). The single hydrocarbon truncation of a glutamine to asparagine ablated GlcCer transport without substantially altering PC selection (Fig. 4c, d). Q^655^ is conserved in all five of the newly established GlcCer P4-ATPases (*S.c.* Dnf1, Dnf2, *S.p.* Dnf2, *H.s.* ATP10A, ATP10D – Figs. 1, 2). These results demonstrate the importance of Q^655^ in GlcCer recognition and suggest it is a conserved determinant of GlcCer translocation. Dnf2^Q655^ precedes the conserved TM4 proline by four residues, and thus will be called the Pro-4 position.

### Conserved requirement for the Pro-4 glutamine and neighboring residues

Based on a Dnf1 homology model (Roland & Graham, 2016b), we mutated residues predicted to surround the Pro-4 position in TM1, 2, 3, and 6, and tested their influence on GlcCer vs. glycerophospholipid transport (Fig. 5a, b, c). Two residues were mutated in TM1, one in TM2, two in TM3, and four in TM6 (Fig. 5d, e). No change in GlcCer coordination was noted for the mutations in TM3 (Fig. 5d, e). In contrast, all positions tested within TM1 and TM2 (mutations at F^261^, L^264^, and L^285^) decreased GlcCer preference (Fig. 5d, e). Four residues were mutated in TM6, revealing three mutations that abrogated GlcCer preference. S^1257^ is predicted to be positioned in the lumenal loop between TM5 and TM6 (Fig. 5b), and Dnf2^S1257Q^ reduced GlcCer and glycerophospholipid transport (Fig. 5e). Residues T^1266^ and L^1270^ may project directly toward Q^655^ along a parallel segment of TM6, and mutating either position impaired GlcCer selection (Fig. 5d, e). T^1266^ mutations in particular strongly affected the GlcCer transport without impairing PC transport (Fig. 5d, e).

**Fig 5.**
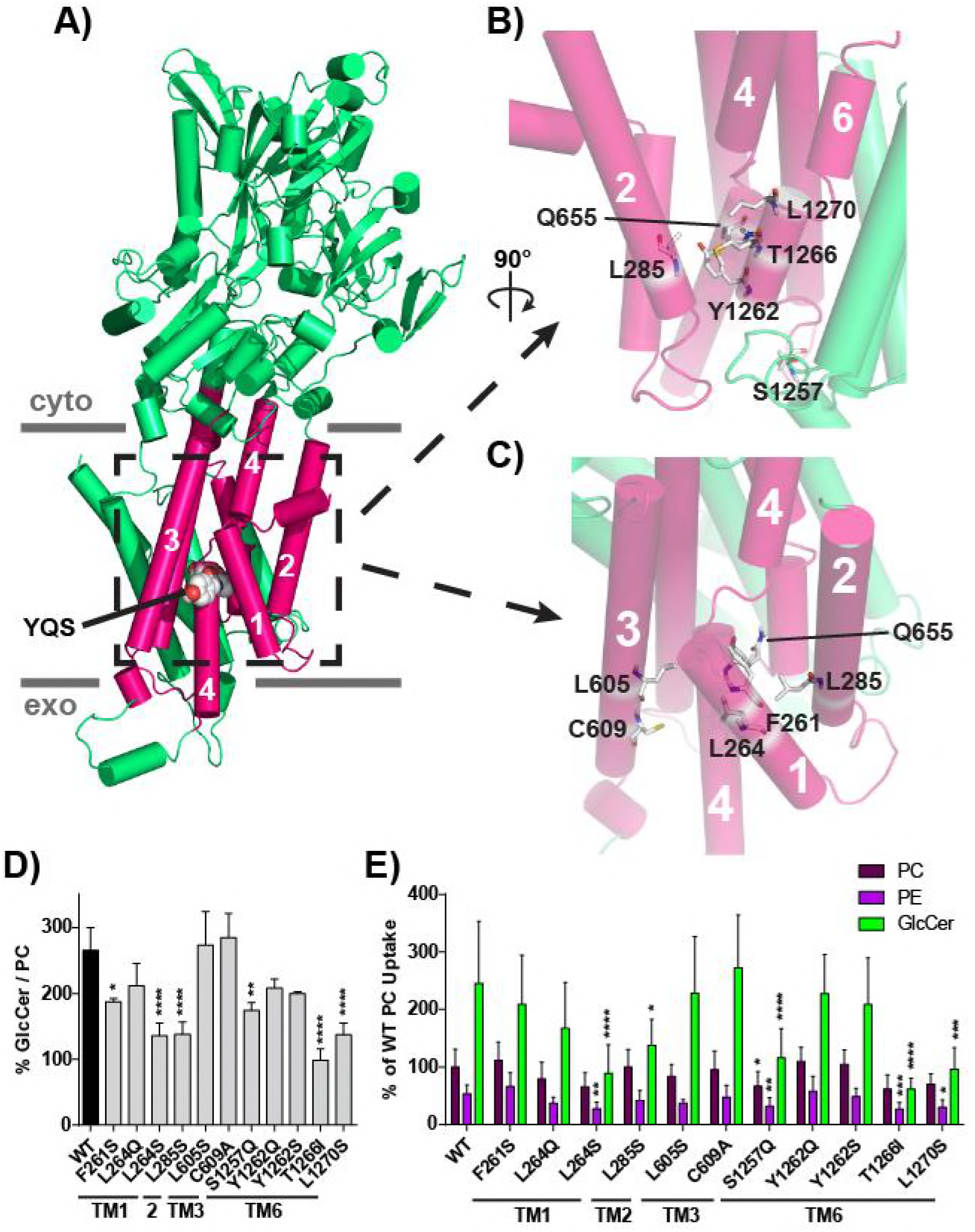
Residues modeled in proximity to the Pro-4 glutamine influence GlcCer transport. **(A)** Homology model of Dnf1 (PDB ID code 3W5D) with TM 1–6 shown as pink cylinders, the rest of the protein colored green, and YQS motif represented in spheres and colored by element; PM boundaries are indicated. **(B)** A 90° rotated and enhanced view of the peri-Q^655^ region formed by TMs 2, 4, and 6. **(C)** An enhanced view of the of TMs 1, 2, 3, and 4 that surround Q^655^. Residues were selected for mutagenesis by identifying positions that were predicated to be planar with the YQS motif, and are shown in sticks and colored by element. (**D-E**) GlcCer preference (**D**) and selection (**E**) were examined for all positions, revealing that TM1, TM2, and TM6 positions were capable of altering GlcCer transport. One position required chemical specificity to alter GlcCer transport (L264). Variance was assessed among data sets using One-way ANOVAs, and comparisons to WT made with Tukey’s post hoc analysis. n ≥ 9, ± SD, * indicates p<0.05, **p<0.01, ***p<0.001, ****p<0.0001.

The T^1266^ residue is not conserved in Dnf1 (Fig S9a, b), therefore we generated a new homology model of Dnf2 to more closely evaluate this region of the enzyme. L^1265^ and T^1266^ are predicted to flank the Q^655^ residue, potentially restricting the rotational flexibility of Q^655^ (Fig S9c). Dnf1 encodes two sequential methionines in the Dnf2 “LT” positions, and a Dnf2^L1265M,T1266M^ construct recapitulated the Dnf1 preference for GlcCer (Fig S9f, g). Replacing the “LT” with the ATP8A2 sequence (“IG” Dnf2^L1265I,T1266G^), an enzyme that is not predicted to transport GlcCer, reduced GlcCer transport without impacting PC coordination (Fig S9f, g). The lack of conservation in this region may suggest that these residues are not primary coordinators of GlcCer binding, but this position may help support a larger substrate binding pocket or translocation pathway.

To test whether the mechanism of GlcCer recognition is conserved, we mutated the Pro-4 and Pro-3 positions in TM4 of ATP10A and ATP10D to assess their influence on PC and GlcCer transport (Fig. 6). The ATP10D^Q381N,V382T^ mutant ablated GlcCer transport (Fig. 6b), similar to its impact in Dnf2. The necessity of the Pro-4 glutamine was confirmed by separating this TM4 motif into single mutants. Again, a simple change of a glutamine-to-asparagine impaired GlcCer transport (Fig. 6c); however, the single ATP10D^V382T^ mutation elicited the unanticipated capacity to enhance GlcCer transport (Fig. 6c). Finally, the ATP10A^Q370N,V371T^ mutant did not impact PC transport (Fig. 6d), thus reinforcing the specific and conserved role that the Pro-4 position plays in GlcCer, but not PC translocation.

**Fig 6.**
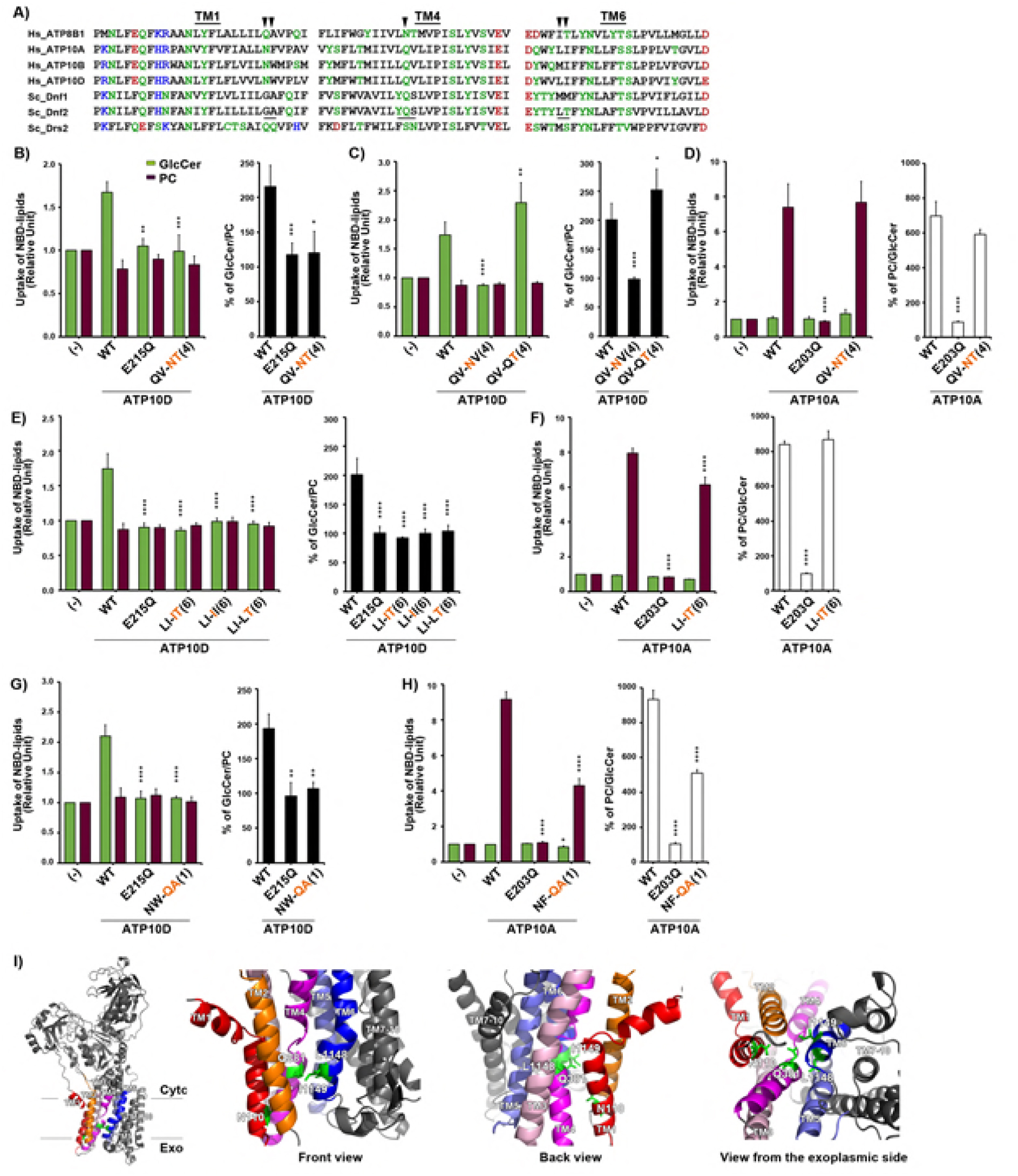
Structure-function analysis of the *H.s.* ATP10D substrate pathway demonstrates primary structural conservation of TM4 Pro-4 position in GlcCer transport. (**A**) Sequence alignments of TM1, TM4, and TM6 of P4-ATPases are shown. Hydrophilic residues are indicated in **green**, **red** (negatively charged), and **blue** (positively charged), and hydrophobic residues are indicated in **black**. Three motifs, which were required for GlcCer preference in Dnf2, were underlined. The arrowheads indicate amino acids which were critical for ATP10D to transport GlcCer. (**B-H**) NBD-lipid uptake was measured in HeLa cells stably expressing C-terminally HA-tagged ATP10A (WT), ATP10D (WT), each mutant (indicated), and parental cells (-); TM is numbered in parenthesis. (**B-D**) The glutamine at TM4 Pro-4 was critical for ATP10D to transport GlcCer but was dispensable for ATP10A to transport PC. (**E, F**) Motif in TM6 was critical for GlcCer transport of ATP10D but was dispensable for PC transport of ATP10A. The experiments of (**E**) panels were performed together with (**C**) panels and thus graphs (-) and WT of (**E**) are equivalent to those of (**C**). Graphs display averages from three independent experiments ± S.D. Fold increase of NBD-lipid uptake compared with parental cells (-) is shown. A one-way ANOVA was performed to assess variance in panels (**B-H**), and comparisons to WT were made with Tukey’s post hoc analysis. * indicates p<0.05, **p<0.01, ***p<0.001, ****p<0.0001. (I) TM1, 2, 3, 4, 5, and 6 are indicated in red, orange, pink, magenta, purple, and blue, respectively, and others are indicated in gray. Critical residues for GlcCer transport are indicated in green sticks.

The TM1 and TM6 residues defined in yeast are not conserved in the mammalian enzymes, yet we tested the influence of these positions on ATP10D and ATP10A GlcCer and PC transport. ATP10D^L1148,I1149^ and ATP10A ^L1123,I1124^ correspond with the TM6 LT motif of Dnf2, and an ATP10D homology model similarly predicted their proximity to the Pro-4 position (Fig. 6i). Both L^1148^ and I^1149^ in TM6 of ATP10D were critical for the transport of GlcCer (Fig. 6e), while the corresponding residues in ATP10A played a minor role in PC transport (Fig. 6f). ATP10D^N110,W111^ and ATP10A^N98,F99^ correspond to the GA motif in TM1 of Dnf2. ATP10D^N110,W111^ substitutions abolished GlcCer transport (Fig. 6g), while ATP10A^N98,F99^ mutations moderately effected PC transport (Fig. 6h). All mutant enzymes were robustly expressed and localized to the plasma membrane (Fig S5a-c), suggesting general enzyme integrity and demonstrating proper trafficking. Collectively, these experiments highlight conserved structural determinants of P4-ATPase-mediated GlcCer transport in TM1, TM4, and TM6. For TM4, a Pro-4 glutamine is critical for GlcCer transport, while positions in TM1 and TM6 are conserved even though the specific amino acids at these sites differ.

## Discussion

Bioactive sphingolipids are linked to numerous cell and physiological processes, including cell death/survival, cell proliferation, senescence, autophagy, migration, differentiation, adhesion and inflammatory responses (Hannun & Obeid, 2018). In addition, dysregulation of sphingolipid metabolism contributes to neurological disease, cancer, metabolic disorder, type 2 diabetes, hepatic steatosis and cardiovascular disease. While ceramides and sphingosines have received the most attention in this context, GlcCer is a central species in the metabolism of sphingolipids. GlcCer synthesis is a major consumer of ceramide in animal cells and its breakdown can produce ceramide and sphingosine, yet GlcCer transport has been a particularly enigmatic topic. Although GlcCer is synthesized in the cytosolic leaflet of the Golgi, an unknown transporter carries it to the Golgi lumen where it becomes the keystone substrate for a majority of more than 2,000 different glycosphingolipids (Fahy, Subramaniam et al., 2009). Mechanisms for GlcCer transfer between serum lipoprotein particles (low-density lipoproteins and very low-density lipoproteins) and cells is also unclear, as is the distribution of GlcCer between leaflets of cellular membranes. Thus, defining the transport mechanisms of this key lipid species will be essential to understanding how sphingolipids contribute to pathogenesis, and how they may be leveraged for the treatment of human disease.

We report here the identification and molecular characterization of a family of GlcCer transporters conserved from yeast to humans. Phylogenetic analyses of *S. cerevisiae*, *S. pombe*, and *H. sapiens* P4-ATPases suggests that *S.c.* Dnf1,2 and *S.p.* Dnf2 cluster with the ATP10 family of human P4-ATPases (Fig. 7). Given the evolutionary distance between these three organisms, we propose that these five P4-ATPases represent a functional clade of glycosphingolipid flippases (Fig. 7). We establish that the yeast enzymes are capable of transporting both NBD-GlcCer and NBD-GalCer, while the human enzymes appear to be specific for NBD-GlcCer. The difference between these two substrates is simply the orientation of a distal hydroxyl, highlighting the exquisite specificity of these enzymes. We have shown in yeast and human P4-ATPases that GlcCer transport is strongly influenced by components of TM1, TM4, and TM6, at positions that are exoplasmic relative to the invariant TM4 proline. These residues do not affect PC transport (Fig S10), leading us to believe that we have not altered essential components required to appropriately fold, traffic, and complete the P4-ATPase catalytic cycle. This mutagenic dissection of yeast and human enzymes leads us to propose that GlcCer is coordinated by a cluster of residues that act cooperatively.

**Fig 7.**
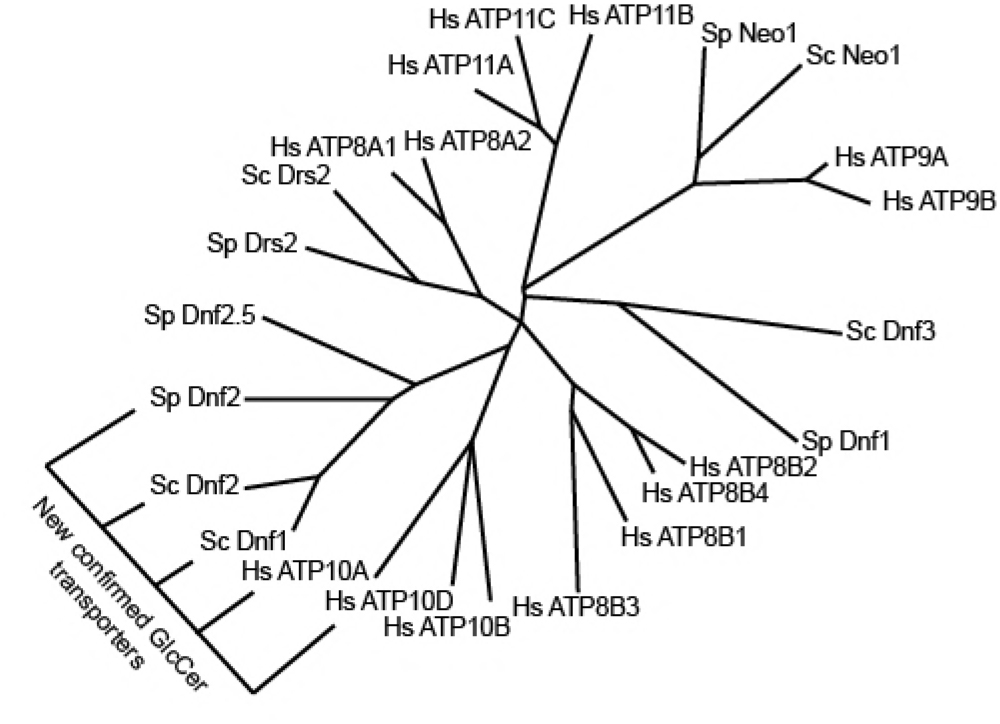
Yeast Dnf1,2 phylogenically clusters with the human ATP10D family members. An unrooted phylogenic tree of *S.cerevisiae*, *S.pombe*, and *H.sapiens* P4-ATPases with branch length indicating character change. A new clade of GlcCer P4-ATPases is indicated. Protein accession numbers and tools used for analysis are found in Experimental Procedures.

There are two prevailing hypotheses that describe the mechanism of P4-ATPase substrate transport, i) a hydrophobic gate model, and ii) a two-gate model (Andersen et al., 2016, Lopez-Marques, Poulsen et al., 2015, Roland & Graham, 2016a). The hydrophobic gate model proposes that substrate translocation is facilitated by the restriction and dilation of a cluster of hydrophobic residues centered around the middle of TM4 (Vestergaard et al., 2014). The two-gate model suggests that “entry” and “exit” gates at the exoplasmic and cytoplasmic membrane interface will bind and transfer substrate along TM4 (Baldridge & Graham, 2013). These models are complementary as they each describe different aspects of the enzymatic process, and indeed, the sequential passage of a substrate through a hydrophobic barrier would facilitate the directionality of P4-ATPase translocation as it transitions from entry to exit gates.

The principal determinant of GlcCer transport is a conserved TM4 glutamine at the Pro-4 position. It was surprising that a Q-to-N mutation was sufficient to ablate GlcCer transport in yeast and human enzymes. Glutamine and asparagine are chemically similar residues, only differing in the extension of their terminal amine (Fig S11a). However, the difference in a hydrocarbon provides additional rotational flexibility to the glutamine side chain, relative to asparagine. Homology models of Dnf2 and ATP10D predict that the Pro-4 glutamine occupies a relatively open pocket within the TM domain of the enzyme (Fig S11b, c). The glutamine side chain was easily sculpted around the β-γ carbon bond within the Dnf2 and ATP10D models, and a 360° rotation around this bond was only restricted by Dnf2^Y1262^, Dnf2^L1265^, and Dnf2^T1266^, or ATP10D^Y1145^, ATP10D^L1148^, and ATP10D^I1149^ (Fig S11b, c). Conversely, the rotation of the terminal amide of asparagine around the β-γ carbon bond was unimpeded (Fig S11d, e). Thus, the flexibility and rotational capacity of the Pro-4 amide may be an important component of GlcCer selection and translocation.

The Pro-4 position was previously proposed to modulate the hydration of a pocket that preceded the hydrophobic gate (Vestergaard et al., 2014). This study defined a conserved isoleucine at the Pro+1 position as the primary component of a hydrophobic gate (Vestergaard et al., 2014). The proximity of these two residues raises the possibility that the Pro-4 glutamine may anchor a pre-hydrophobic gate GlcCer binding pocket, similar to that proposed for phosphatidylserine coordination in ATP8A2 (Vestergaard et al., 2014). However, it is clear that not all P4-ATPase substrates require these residues for transport, and possibly use an alternative substrate pathway. The separation of function mutations in yeast Dnf1 and Dnf2 demonstrate that the Pro-4 and its surrounding residues do not impact PC or PE transport, and mutagenesis data with human ATP10A and ATP10D supports this separation of function. These results demonstrate that PC and GlcCer are coordinated differently, and may be transported by independent pathways.

Fungi are heterotrophic eukaryotic organisms that will absorb materials from their environment to facilitate growth. Most yeast produce GlcCer, but both *S. pombe* and *S. cerevisiae* have independently lost the GlcCer synthase enzyme (Leipelt, Warnecke et al., 2001). Therefore, the GlcCer transport activity in these organisms may have evolved to scavenge this lipid from plant material that yeast grow upon (Jiang, Wang et al., 2017, Luginbuehl, Menard et al., 2017). This possibility would explain why Dnf1 and Dnf2 are intimately linked to sphingolipid homeostatic pathways (Roelants et al., 2010), even though *S. cerevisiae* and *S. pombe* do not endogenously produce these glycolipids. The translocation of this substrate could be leveraged to change membrane properties, as GlcCer supplementation has been shown to increase alkaline resistance in yeast (Sawada, Sato et al., 2015). Alternatively, imported GlcCer could be catabolized and its backbone, N-linked acyl chain, and/or headgroup repurposed.

Membrane asymmetry is the difference in lipid composition between the two leaflets of the membrane, and is a critical property of the plasma membrane. Simple alterations of this asymmetric structure can elicit substantial changes in biology, influencing micro and macro-processes such as vesicle budding (Chen, Ingram et al., 1999, Gall, Geething et al., 2002, Liu, Surendhran et al., 2008, Xu, Baldridge et al., 2013), apoptosis (Bratton, Fadok et al., 1997, Fadok, Bratton et al., 1998, Uchida, Emoto et al., 1998), phagocytosis (Fadok, Bratton et al., 2000, Hoffmann, deCathelineau et al., 2001, Yoshida, Kawane et al., 2005), as well as many others. The majority of research has thus far examined the cellular and physiologic impact of glycerophospholipid asymmetry, yet our discovery of two endogenous human GlcCer flippases raises important questions about the impact of sphingolipid asymmetry. For example, it is known that there are three human glucocerebrosidases, enzymes that cleave the glucose headgroup from the ceramide backbone. Glucocerebrosidase 1 is the best understood, and mutations in the *GBA1* gene lead to Gaucher’s disease and Parkinson’s disease (Platt, 2014, Sidransky & Lopez, 2012, Vitner & Futerman, 2013). Glucocerebrosidase 1 localizes to the lumen of the lysosome, yet glucocerebrosidase 2 and 3 localize to the cytosol (Astudillo, Therville et al., 2016). It is unclear where glucocerebrosidase 2 and 3 obtain their cytosolic GlcCer substrates, and what role they perform in sphingolipid homeostasis. ATP10A and ATP10D are importers of GlcCer, and therefore may be providing cytosolic substrates for metabolism or signaling. Thus, we hypothesize that the transport activity of ATP10A and ATP10D toward GlcCer may be an important component of sphingolipid homeostasis and lipid metabolism in mammals.

### Experimental Procedures

#### Reagents

All yeast culture reagents were purchased from Sigma Aldrich, as well as NBD-hexanoic acid. All other lipids used in this study were purchased from Avanti Polar Lipids (Fig S2). TLC materials and solvents used for lipid extraction were purchased from Fisher Scientific. *Strains and Culture. S.pombe* were cultured on YES media, and *S.cerevisiae* were grown on synthetic defined (SD) media unless examined in parallel with *S.pombe*. If examined in parallel with *S.pombe*, *S.cerevisiae* were grown and analyzed on YPD. All yeast and human cell lines used in the study are listed in Table S1, and plasmids are listed in Table S2.

#### Establishment of HeLa stable cell lines, antibodies and immunofluorescence analysis

HeLa cells were cultured as described previously (Takatsu et al., 2011). HeLa cells expressing each C-terminally HA-tagged ATP8B4 and ATP10D(E215Q) were established as described previously (Naito et al., 2015, Takatsu et al., 2014). Sources of antibodies used in the present study were as follows: monoclonal rabbit anti-ATP1A1 (EP1845Y), Abcam; monoclonal rat anti-HA (3F10), Roche Applied Science; Alexa Fluor–conjugated secondary antibodies, Molecular Probes; Cy3-, and horseradish peroxidase–conjugated secondary antibodies, Jackson ImmunoResearch Laboratories. Immunostaining was performed as described previously (Shin, Morinaga et al., 2004) and visualized using an Axiovert 200MAT microscope (Carl Zeiss, Thornwood, NY)

#### Lipid Administration and Flow Cytometry (Yeast cells)

NBD-lipid administration and analysis were performed as described previously (Roland & Graham, 2016b). Briefly, overnight yeast cultures were subcultured to 0.15 OD_600_/ml, cultured to mid-log phase, and 0.5 ml of cells were collected per sample. The designated NBD-lipid was solubilized in 100% ethanol and added to the designated ice cold growth media at a final concentration of 2 μg/ml with final ethanol volumes ≤0.5%. Cells were suspended with the designated media + NBD-lipids and incubated on ice for 30 min, unless otherwise stated (kinetics analyses and TLC examinations). After 30 min, the cells were washed twice with ice cold SA (SD + 2% [wt/vol] sorbitol + 20 mM NaN_3_) supplemented with 4% (wt/vol) fatty acid free bovine serum albumin. All cells regardless of original media growth were washed with ice cold SD containing either dropout or complete amino acid supplements, respective the strain/transformant requirements. Once washed with chilled 4% BSA-SA, cells were washed with ice cold SA, resuspended in chilled SA containing 5 μM propidium iodide (PI), and analyzed immediately. The sequence of sample preparation and processing was varied in each experiment to mitigate potential positional experimental bias. Additional detail is provided in the Supplemental Methods.

#### NBD-lipid Administration and Flow Cytometry (Human cells)

The incorporation of NBD-lipids was analyzed by flow cytometry as described (Takatsu et al., 2014) with some modifications. In brief, HeLa cells (in 24-well plate) were washed and equilibrated at 15°C for 15 min in 500 μl of Hank’s balanced salt solution (pH 7.4) containing 1 g/l glucose (HBSS-glucose). The buffer was replaced with 1 μM NBD-lipid in HBSS-glucose and cells were further incubated at 15°C. After incubation for indicated times, the buffer was replaced with ice-cold PBS(-) containing 2.5% (w/v) fatty acid–free BSA (Wako), 5 mM EDTA, and 0.5 μg/ml of propidium iodide (Nacalai Tesque) and cells were incubated on ice for 30 min. The detached cells (more than 10^4^ cells / sample) were analyzed with a FACSCalibur (BD Biosciences) to measure fluorescence of NBD-lipids incorporated and translocated into the cytoplasmic leaflet of the plasma membrane. Graphs for NBD-lipid flippase activities are expressed as the averages of three independent experiments ± SD. settings.

#### Protein homology modeling

All structural images were generated using PyMol. The Dnf1 homology model was previously published (Roland & Graham, 2016b). Dnf2 and ATP10D homology models used for sculpting were generated using the intensive Phyre^2^ modeling (Kelley, Mezulis et al., 2015). The Phyre2 intensive modeling process represents an unbiased approach. First, homologous templates are surveyed, crude backbone models are generated, and ranked by coverage and confidence; then loop fragments are modeled in 2-15 residue chunks. A protein folding simulator uses a heuristic process to synthesize multiple templates, and leverages *ab inito* strategies for regions without predictions. Finally, side chains are placed using a rotamer library, optimized to avoid steric clashes. Unlike the threaded modeling approach, the intensive modeling process facilitates subsequent model manipulation. The Dnf2 model was generated from the synthesis of PDB IDs: 1MHS(Kuhlbrandt, Zeelen et al., 2002), 4WIT(Brunner, Lim et al., 2014), 3B9B(Olesen, Picard et al., 2007), 3IXZ(Abe, Tani et al., 2009), 2ZXE(Shinoda, Ogawa et al., 2009), 3B8C(Focht, Croll et al., 2017), 3B8E(Morth, Pedersen et al., 2007); while the ATP10D intensive model used PDB IDs: 1MHS(Kuhlbrandt et al., 2002), 3B9B(Olesen et al., 2007), 3IXZ(Abe et al., 2009), 2ZXE(Shinoda et al., 2009), 3B8C(Focht et al., 2017), and 3B8E(Morth et al., 2007). Residue sculpting within these models was manually performed via PyMol, with residue shells and cushions set at 6 Å. The ATP10D model used in Fig. 6 was generated by threading the sequence over the structure of the Na/K pump (PDB ID: 2ZXE) (Shinoda et al., 2009) using Phyre^2^ (Kelley et al., 2015).

#### Data Analysis

Yeast substrate uptake data analyses were performed as previously outlined (Roland & Graham, 2016b). Briefly, three independent clonal transformants were selected for each yeast experiment and examined in parallel per sample. When KO strains were examined, three clonal strain cultures were assessed in parallel. When reported as A.F.U., raw fluorescence data is presented. When normalized to WT NBD-PC transport, substrate uptake in *dnf1,2∆* vector controls are subtracted from the experimental group, and the data are normalized relative to WT PC uptake, which is set at 100%.

Median fluorescence values were averaged from the respective experiments and substrate preference was determined by taking ratios of clonal replicates.

## Author contributions

B.P.R. and T.R.G. conceived of the project; B.P.R., T.N., C.A.-Y., R.J.Y., H.-W.S. and T.R.G. designed the experiments; B.P.R., T.N., J.T.B., R.J.Y., H.T., and C.A.-Y. performed the experiments; B.P.R., T.N., J.T.B., H.-W.S., and T.R.G. analyzed the data; and B.P.R., H.-W.S., and T.R.G. wrote the manuscript.

## Acknowledgements

We thank the laboratory of Kathleen L. Gould (Vanderbilt University) for their gift of the *S.pombe* strains. We thank Toshio Kitamura (University of Tokyo) and Hiroyuki Miyoshi (RIKEN BioResource Center) for providing plasmids for retroviral infection. Yeast flow cytometry experiments were performed with the Vanderbilt Medical Center Flow Cytometry Shared Resource, supported by National Institutes of Health (NIH) grants to the Vanderbilt Ingram Cancer Center (P30-CA68485) and Vanderbilt Digestive Disease Research Center (P30-DK058404). This work was supported by NIH grants R01-GM107978 (to T.R.G.) and F32-GM116310 (to B.P.R.), JSPS KAKENHI Grant Numbers JP16H00764, and JP17H03655, and grant from the Japan Foundation for Applied Enzymology (to H.-W.S) and Grant-in-Aid for JSPS research fellow JP16J05381 (to T.N.).

## References

Abe K, Tani K, Nishizawa T, Fujiyoshi Y (2009) Inter-subunit interaction of gastric H+,K+-ATPase prevents reverse reaction of the transport cycle. EMBO J 28: 1637-43

Andersen JP, Vestergaard AL, Mikkelsen SA, Mogensen LS, Chalat M, Molday RS (2016) P4-ATPases as Phospholipid Flippases-Structure, Function, and Enigmas. Front Physiol 7: 275

Astudillo L, Therville N, Colacios C, Segui B, Andrieu-Abadie N, Levade T (2016) Glucosylceramidases and malignancies in mammals. Biochimie 125: 267-80

Baldridge RD, Graham TR (2012) Identification of residues defining phospholipid flippase substrate specificity of type IV P-type ATPases. Proc Natl Acad Sci U S A 109: E290-8

Baldridge RD, Graham TR (2013) Two-gate mechanism for phospholipid selection and transport by type IV P-type ATPases. Proc Natl Acad Sci U S A 110: E358-67

Bratton DL, Fadok VA, Richter DA, Kailey JM, Guthrie LA, Henson PM (1997) Appearance of phosphatidylserine on apoptotic cells requires calcium-mediated nonspecific flip-flop and is enhanced by loss of the aminophospholipid translocase. J Biol Chem 272: 26159-65

Brunner JD, Lim NK, Schenck S, Duerst A, Dutzler R (2014) X-ray structure of a calcium-activated TMEM16 lipid scramblase. Nature 516: 207-12

Bryde S, Hennrich H, Verhulst PM, Devaux PF, Lenoir G, Holthuis JC (2010) CDC50 proteins are critical components of the human class-1 P4-ATPase transport machinery. J Biol Chem 285: 40562-72

Bull LN, van Eijk MJ, Pawlikowska L, DeYoung JA, Juijn JA, Liao M, Klomp LW, Lomri N, Berger R, Scharschmidt BF, Knisely AS, Houwen RH, Freimer NB (1998) A gene encoding a P-type ATPase mutated in two forms of hereditary cholestasis. Nat Genet 18: 219-24

Chen CY, Ingram MF, Rosal PH, Graham TR (1999) Role for Drs2p, a P-type ATPase and potential aminophospholipid translocase, in yeast late Golgi function. J Cell Biol 147: 1223-36

Coleman JA, Zhu X, Djajadi HR, Molday LL, Smith RS, Libby RT, John SW, Molday RS (2014) Phospholipid flippase ATP8A2 is required for normal visual and auditory function and photoreceptor and spiral ganglion cell survival. J Cell Sci 127: 1138-49

Dawson G, Kruski AW, Scanu AM (1976) Distribution of glycosphingolipids in the serum lipoproteins of normal human subjects and patients with hypo-and hyperlipidemias. J Lipid Res 17: 125-31

Dhar M, Hauser L, Johnson D (2002) An aminophospholipid translocase associated with body fat and type 2 diabetes phenotypes. Obes Res 10: 695-702

Dhar MS, Sommardahl CS, Kirkland T, Nelson S, Donnell R, Johnson DK, Castellani LW (2004) Mice heterozygous for Atp10c, a putative amphipath, represent a novel model of obesity and type 2 diabetes. J Nutr 134: 799-805

Fadok VA, Bratton DL, Frasch SC, Warner ML, Henson PM (1998) The role of phosphatidylserine in recognition of apoptotic cells by phagocytes. Cell Death Differ 5: 551-62

Fadok VA, Bratton DL, Rose DM, Pearson A, Ezekewitz RA, Henson PM (2000) A receptor for phosphatidylserine-specific clearance of apoptotic cells. Nature 405: 85-90

Fahy E, Subramaniam S, Murphy RC, Nishijima M, Raetz CR, Shimizu T, Spener F, van Meer G, Wakelam MJ, Dennis EA (2009) Update of the LIPID MAPS comprehensive classification system for lipids. J Lipid Res 50 Suppl: S9-14

Flamant S, Pescher P, Lemercier B, Clement-Ziza M, Kepes F, Fellous M, Milon G, Marchal G, Besmond C (2003) Characterization of a putative type IV aminophospholipid transporter P-type ATPase. Mamm Genome 14: 21-30

Focht D, Croll TI, Pedersen BP, Nissen P (2017) Improved Model of Proton Pump Crystal Structure Obtained by Interactive Molecular Dynamics Flexible Fitting Expands the Mechanistic Model for Proton Translocation in P-Type ATPases. Front Physiol 8: 202

Folmer DE, Elferink RP, Paulusma CC (2009) P4 ATPases - lipid flippases and their role in disease. Biochim Biophys Acta 1791: 628-35

Gall WE, Geething NC, Hua Z, Ingram MF, Liu K, Chen SI, Graham TR (2002) Drs2p-dependent formation of exocytic clathrin-coated vesicles in vivo. Curr Biol 12: 1623-7

Gantzel RH, Mogensen LS, Mikkelsen SA, Vilsen B, Molday RS, Vestergaard AL, Andersen JP (2017) Disease mutations reveal residues critical to the interaction of P4-ATPases with lipid substrates. Sci Rep 7: 10418

Hannun YA, Obeid LM (2018) Sphingolipids and their metabolism in physiology and disease. Nat Rev Mol Cell Biol 19: 175-191

Hicks AA, Pramstaller PP, Johansson A, Vitart V, Rudan I, Ugocsai P, Aulchenko Y, Franklin CS, Liebisch G, Erdmann J, Jonasson I, Zorkoltseva IV, Pattaro C, Hayward C, Isaacs A, Hengstenberg C, Campbell S, Gnewuch C, Janssens AC, Kirichenko AV et al. (2009) Genetic determinants of circulating sphingolipid concentrations in European populations. PLoS Genet 5: e1000672

Hoffmann PR, deCathelineau AM, Ogden CA, Leverrier Y, Bratton DL, Daleke DL, Ridley AJ, Fadok VA, Henson PM (2001) Phosphatidylserine (PS) induces PS receptor-mediated macropinocytosis and promotes clearance of apoptotic cells. J Cell Biol 155: 649-59

Irvin MR, Wineinger NE, Rice TK, Pajewski NM, Kabagambe EK, Gu CC, Pankow J, North KE, Wilk JB, Freedman BI, Franceschini N, Broeckel U, Tiwari HK, Arnett DK (2011) Genome-wide detection of allele specific copy number variation associated with insulin resistance in African Americans from the HyperGEN study. PLoS One 6: e24052

Jensen MS, Costa SR, Duelli AS, Andersen PA, Poulsen LR, Stanchev LD, Gourdon P, Palmgren M, Gunther Pomorski T, Lopez-Marques RL (2017) Phospholipid flipping involves a central cavity in P4 ATPases. Sci Rep 7: 17621

Jiang Y, Wang W, Xie Q, Liu N, Liu L, Wang D, Zhang X, Yang C, Chen X, Tang D, Wang E (2017) Plants transfer lipids to sustain colonization by mutualistic mycorrhizal and parasitic fungi. Science 356: 1172-1175

Kelley LA, Mezulis S, Yates CM, Wass MN, Sternberg MJ (2015) The Phyre2 web portal for protein modeling, prediction and analysis. Nat Protoc 10: 845-58

Kengia JT, Ko KC, Ikeda S, Hiraishi A, Mieno-Naka M, Arai T, Sato N, Muramatsu M, Sawabe M (2013) A gene variant in the Atp10d gene associates with atherosclerotic indices in Japanese elderly population. Atherosclerosis 231: 158-62

Kuhlbrandt W, Zeelen J, Dietrich J (2002) Structure, mechanism, and regulation of the Neurospora plasma membrane H+-ATPase. Science 297: 1692-6

Leipelt M, Warnecke D, Zahringer U, Ott C, Muller F, Hube B, Heinz E (2001) Glucosylceramide synthases, a gene family responsible for the biosynthesis of glucosphingolipids in animals, plants, and fungi. J Biol Chem 276: 33621-9

Liu K, Surendhran K, Nothwehr SF, Graham TR (2008) P4-ATPase requirement for AP-1/clathrin function in protein transport from the trans-Golgi network and early endosomes. Mol Biol Cell 19: 3526-35

Lopez-Marques RL, Poulsen LR, Bailly A, Geisler M, Pomorski TG, Palmgren MG (2015) Structure and mechanism of ATP-dependent phospholipid transporters. Biochim Biophys Acta 1850: 461-75

Luginbuehl LH, Menard GN, Kurup S, Van Erp H, Radhakrishnan GV, Breakspear A, Oldroyd GED, Eastmond PJ (2017) Fatty acids in arbuscular mycorrhizal fungi are synthesized by the host plant. Science 356: 1175-1178

Morth JP, Pedersen BP, Toustrup-Jensen MS, Sorensen TL, Petersen J, Andersen JP, Vilsen B, Nissen P (2007) Crystal structure of the sodium-potassium pump. Nature 450: 1043-9

Naito T, Takatsu H, Miyano R, Takada N, Nakayama K, Shin HW (2015) Phospholipid Flippase ATP10A Translocates Phosphatidylcholine and Is Involved in Plasma Membrane Dynamics. J Biol Chem 290: 15004-17

Natarajan P, Wang J, Hua Z, Graham TR (2004) Drs2p-coupled aminophospholipid translocase activity in yeast Golgi membranes and relationship to in vivo function. Proc Natl Acad Sci U S A 101: 10614-9

Olesen C, Picard M, Winther AM, Gyrup C, Morth JP, Oxvig C, Moller JV, Nissen P (2007) The structural basis of calcium transport by the calcium pump. Nature 450: 1036-42

Onat OE, Gulsuner S, Bilguvar K, Nazli Basak A, Topaloglu H, Tan M, Tan U, Gunel M, Ozcelik T (2013) Missense mutation in the ATPase, aminophospholipid transporter protein ATP8A2 is associated with cerebellar atrophy and quadrupedal locomotion. Eur J Hum Genet 21: 281-5

Platt FM (2014) Sphingolipid lysosomal storage disorders. Nature 510: 68-75

Pomorski T, Lombardi R, Riezman H, Devaux PF, van Meer G, Holthuis JC (2003) Drs2p-related P-type ATPases Dnf1p and Dnf2p are required for phospholipid translocation across the yeast plasma membrane and serve a role in endocytosis. Mol Biol Cell 14: 1240-54

Roelants FM, Baltz AG, Trott AE, Fereres S, Thorner J (2010) A protein kinase network regulates the function of aminophospholipid flippases. Proc Natl Acad Sci U S A 107: 34-9

Roland BP, Graham TR (2016a) Decoding P4-ATPase substrate interactions. Crit Rev Biochem Mol Biol 51: 513-527

Roland BP, Graham TR (2016b) Directed evolution of a sphingomyelin flippase reveals mechanism of substrate backbone discrimination by a P4-ATPase. Proc Natl Acad Sci U S A 113: E4460-6

Sawada K, Sato T, Hamajima H, Jayakody LN, Hirata M, Yamashiro M, Tajima M, Mitsutake S, Nagao K, Tsuge K, Abe F, Hanada K, Kitagaki H (2015) Glucosylceramide Contained in Koji Mold-Cultured Cereal Confers Membrane and Flavor Modification and Stress Tolerance to Saccharomyces cerevisiae during Coculture Fermentation. Appl Environ Microbiol 81: 3688-98

Shin HW, Morinaga N, Noda M, Nakayama K (2004) BIG2, a guanine nucleotide exchange factor for ADP-ribosylation factors: its localization to recycling endosomes and implication in the endosome integrity. Mol Biol Cell 15: 5283-94

Shinoda T, Ogawa H, Cornelius F, Toyoshima C (2009) Crystal structure of the sodium-potassium pump at 2.4 A resolution. Nature 459: 446-50

Sidransky E, Lopez G (2012) The link between the GBA gene and parkinsonism. Lancet Neurol 11: 986-98

Sigruener A, Wolfrum C, Boettcher A, Kopf T, Liebisch G, Orso E, Schmitz G (2017) Lipidomic and metabolic changes in the P4-type ATPase ATP10D deficient C57BL/6J wild type mice upon rescue of ATP10D function. PLoS One 12: e0178368

Stone A, Chau C, Eaton C, Foran E, Kapur M, Prevatt E, Belkin N, Kerr D, Kohlin T, Williamson P (2012) Biochemical characterization of P4-ATPase mutations identified in patients with progressive familial intrahepatic cholestasis. J Biol Chem 287: 41139-51

Surwit RS, Feinglos MN, Rodin J, Sutherland A, Petro AE, Opara EC, Kuhn CM, Rebuffe-Scrive M (1995) Differential effects of fat and sucrose on the development of obesity and diabetes in C57BL/6J and A/J mice. Metabolism 44: 645-51

Takada N, Naito T, Inoue T, Nakayama K, Takatsu H, Shin HW (2018) Phospholipid-flipping activity of P4-ATPase drives membrane curvature. EMBO J 37

Takatsu H, Baba K, Shima T, Umino H, Kato U, Umeda M, Nakayama K, Shin HW (2011) ATP9B, a P4-ATPase (a putative aminophospholipid translocase), localizes to the trans-Golgi network in a CDC50 protein-independent manner. J Biol Chem 286: 38159-67

Takatsu H, Tanaka G, Segawa K, Suzuki J, Nagata S, Nakayama K, Shin HW (2014) Phospholipid flippase activities and substrate specificities of human type IV P-type ATPases localized to the plasma membrane. J Biol Chem 289: 33543-56

Uchida K, Emoto K, Daleke DL, Inoue K, Umeda M (1998) Induction of apoptosis by phosphatidylserine. J Biochem 123: 1073-8

van der Velden LM, Wichers CG, van Breevoort AE, Coleman JA, Molday RS, Berger R, Klomp LW, van de Graaf SF (2010) Heteromeric interactions required for abundance and subcellular localization of human CDC50 proteins and class 1 P4-ATPases. J Biol Chem 285: 40088-96

Vance DE, Vance JE (2008) Biochemistry of lipids, lipoproteins and membranes. Elsevier, Amsterdam; Boston

Vestergaard AL, Coleman JA, Lemmin T, Mikkelsen SA, Molday LL, Vilsen B, Molday RS, Dal Peraro M, Andersen JP (2014) Critical roles of isoleucine-364 and adjacent residues in a hydrophobic gate control of phospholipid transport by the mammalian P4-ATPase ATP8A2. Proc Natl Acad Sci U S A 111: E1334-43

Vitner EB, Futerman AH (2013) Neuronal forms of Gaucher disease. Handb Exp Pharmacol: 405-19

Xu P, Baldridge RD, Chi RJ, Burd CG, Graham TR (2013) Phosphatidylserine flipping enhances membrane curvature and negative charge required for vesicular transport. J Cell Biol 202: 875-86

Yoshida H, Kawane K, Koike M, Mori Y, Uchiyama Y, Nagata S (2005) Phosphatidylserine-dependent engulfment by macrophages of nuclei from erythroid precursor cells. Nature 437: 754-8

Zhu X, Libby RT, de Vries WN, Smith RS, Wright DL, Bronson RT, Seburn KL, John SW (2012) Mutations in a P-type ATPase gene cause axonal degeneration. PLoS Genet 8: e1002853

